# The Monophyly of *Nycteria* and *Polychromophilus* Parasites: A Missing Piece in the Evolution of Malaria and Other Haemosporida

**DOI:** 10.64898/2026.04.07.717123

**Authors:** M. Andreína Pacheco, Juliane Schaer, Oskar Werb, Beatriz Mello, Ananías A. Escalante

**Affiliations:** Biology Department/Institute of Genomics and Evolutionary Medicine (iGEM), Temple University, Philadelphia, PA 19122-1801, USA; Department of Molecular Parasitology, Institute of Biology, Humboldt University, Philippstr. 13, 10115, Berlin, Germany; Department of Biology, Muni University, P.O. Box 725, Arua, Uganda; Department of Biological Sciences, Macquarie University, North Ryde, 2109, NSW, Australia; Department of Genetics, Federal University of Rio de Janeiro, Rio de Janeiro, Brazil

**Keywords:** Chiroptera, Haemosporida, mitochondrial genomes, molecular dating, molecular phylogenetics, nuclear genes

## Abstract

Haemosporida is a diverse order of vector-borne apicomplexan parasites infecting terrestrial vertebrates worldwide, including humans, but the evolutionary relationships among its genera remain unresolved. The phylogenetic placement of two bat-restricted genera, *Nycteria* and *Polychromophilus*, both of which lack erythrocytic schizogony, has varied across studies depending on taxon sampling and marker choice. To address this problem, an expanded dataset of near-complete mitochondrial (mtDNA) genomes together with nine nuclear loci were analyzed. Phylogenetic analyses of mtDNA recovered *Nycteria* and *Polychromophilus* as a strongly supported monophyletic clade. In contrast, analyses based only on the three mitochondrial coding genes (CDS) or a reduced nuclear dataset failed to recover their monophyly and showed low support and extensive topological conflict at deeper nodes. These results indicate that near-complete mitochondrial genomes recover phylogenetic signal that is not captured by reduced mitochondrial coding sequences or partial nuclear datasets. Molecular dating analyses further showed that divergence estimates for a putative *Nycteria*–*Polychromophilus* clade are compatible with the proposed times for bats diversification, and consistent with the broader haemosporidian timescale. When the *Nycteria–Polychromophilus* clade was incorporated as a calibration prior, divergence-time estimates became more precise without altering the overall evolutionary timeframe. Substantial mitochondrial gene-order rearrangements in a distinct *Nycteria* lineage were confirmed, highlighting structural divergence within this bat-associated group. In addition, heterogeneity in rates across mtDNA haemosporidian lineages was observed. Together, these findings support the existence of a distinct bat-associated clade whose deeper placement and evolutionary significance should be tested with broader phylogenomic sampling.

**Author Summary:** Malaria parasites belong to a diverse group of organisms that infect many kinds of vertebrates, including birds, reptiles, and mammals (such as humans). Understanding how these parasites are related to each other is important for explaining how key biological traits have evolved. However, the relationships among major groups of haemosporidian parasites, including malaria parasites, remain unclear, particularly for those infecting bats. In this study, we focused on two groups of bat parasites, *Nycteria* and *Polychromophilus*, which share unusual biological features. The inferred evolutionary relationships of these two genera to other haemosporidians have been inconsistent across previous studies. By analyzing near-complete mitochondrial genomes, we found strong evidence that these two groups descended from a common evolutionary ancestor. In contrast, smaller datasets including nuclear genes failed to recover this relationship and produced conflicting results, suggesting that they lack sufficient information to resolve deep evolutionary relationships. We also found that this bat-associated lineage likely originated around the same time as early bats. In addition, we identified structural changes in the mitochondrial genome of one lineage, highlighting its evolutionary distinctiveness. Together, our results suggest that bats host a unique group of malaria parasites and demonstrate that more complete genetic data are essential for resolving their evolutionary history.

## Introduction

The order Haemosporida (Phylum Apicomplexa) encompasses a diverse radiation of parasite species across terrestrial ecosystems. These parasites are characterized by a complex, heteroxenous life cycle involving Diptera vectors (true flies) and a wide range of vertebrate hosts. Despite their global distribution and significance to human and animal health, the evolutionary relationships within this order remain unresolved [1]. Determining these relationships is critical not only for a reproducible taxonomy but also for understanding the origin and genetic architecture of phenotypes of biomedical importance [1,2].

Classical haemosporidian taxonomy relied on morphological and life-history traits, but early molecular studies demonstrated that these characteristics showed convergence in the phylogeny [3,4]. Mitochondrial (mtDNA) markers, particularly fragments of the cytochrome b gene (*cytb*) [3], greatly expanded taxon sampling but often yielded conflicting phylogenetic hypotheses, likely because of the limited number of informative sites [5]. More recent studies based on near-complete mitochondrial genomes and/or multilocus nuclear genes have not fully resolved the relationships among genera within the order [2,6–12]. Collectively, these analyses indicate that there is not a single origin of Haemosporida in mammals and that host-switching among vertebrate hosts is pervasive [7,13–17].

Among the mammalian haemosporidians, the genera *Nycteria* and *Polychromophilus* occupy a distinct biological niche: both are restricted to bats and lack erythrocytic schizogony [13,18]. Despite these shared characteristics, their phylogenetic relationships remain unstable across molecular phylogenetic studies. Analyses based on a concatenated alignment of partial mitochondrial coding sequences, *clpc* gene, and nuclear loci recovered these genera to be polyphyletic or, alternatively, associated with rodent- or primate-infecting *Plasmodium*, often with weak support [13,19]. Expanded sampling led to the proposal that chiropterans may have played a central role in the radiation of mammalian malaria parasites [7,10]. Clarifying the evolutionary placement of *Nycteria* and *Polychromophilus* is therefore essential for reconstructing patterns of host switching and trait evolution within the order.

Here, an expanded dataset of near-complete mitochondrial genomes together with nuclear markers from *Nycteria* and *Polychromophilus* within Haemosporida was analyzed. Our results show that *Nycteria* and *Polychromophilus* form a well-supported monophyletic lineage in mitochondrial genome analyses, providing the first evidence for this relationship. In contrast, reduced mitochondrial datasets and the currently available nuclear data yield conflicting placements with low support. These results indicate that reduced datasets remain insufficient to consistently resolve the relationships among these taxa [9]. Together, our findings refine the mitochondrial framework for understanding haemosporidian diversification and highlight the need for broader phylogenomic sampling to resolve deep relationships within the order.

## Results

### Organization of the *Nycteria* and *Polychromophilus* Mitochondrial Genome (mtDNA)

A total of 162 mitochondrial (mtDNA) sequences were curated, annotated following Feagin et al. [20], and analyzed, including the newly generated complete mtDNA genomes from the genera *Nycteria* sp. (n=2) and *Polychromophilus* sp. (n=1). Conserved synteny was observed in *Nycteria* sp. (NW3348), *Polychromophilus* sp. (NW3328), and the remaining 159 haemosporidian sequences analyzed. All genomes exhibited the extreme rRNA fragmentation described for *Plasmodium falciparum* [20]. In contrast, the mtDNA genome of *Nycteria* sp. (NW2704) showed a genomic reorganization consistent with that reported for *Nycteria medusiformis* (accession number KX090645) and a related *Nycteria* species (KX090646) [21].

Analysis of putative initiation codons revealed that the *cox3* gene uses alternative TTT (Phe) and TTC (Phe) start sites in *Nycteria* and *Polychromophilus* species, respectively. In the *cox1* gene, the alternative initiation codon TTT (Phe) was conserved across both genera, while the *cytb* gene used the standard ATG (Met) start codon. Termination was mediated by TAA stop codons in *cox1* and *cox3* genes for both genera, while the *cytb* gene utilized either TAA or TAG.

### Phylogenetic analyses: Nuclear versus mitochondrial genomes

#### Mitochondrial phylogeny

Two mitochondrial phylogenetic hypotheses, one based on near-complete mtDNA genomes and a second restricted to the coding sequences from the three mitochondrial genes (CDS), were estimated using a Bayesian Inference (BI) method implemented in MrBayes v3.2.7 with the default priors [22] and a Maximum Likelihood (ML) method executed in IQ-TREE v2.3.1 [23]. For the near-complete mtDNA dataset (159 sequences and 5,052 bp excluding gaps), BI and ML estimates show congruent topologies with high support across most nodes (Fig. 1). Thus, only the BI topology is shown, with posterior probabilities (PP) and ML bootstrap values (B) reported for comparison (Fig 1). Differences between methods were limited to a small number of nodes, most of them within lizard parasite clades.

**Fig 1.**
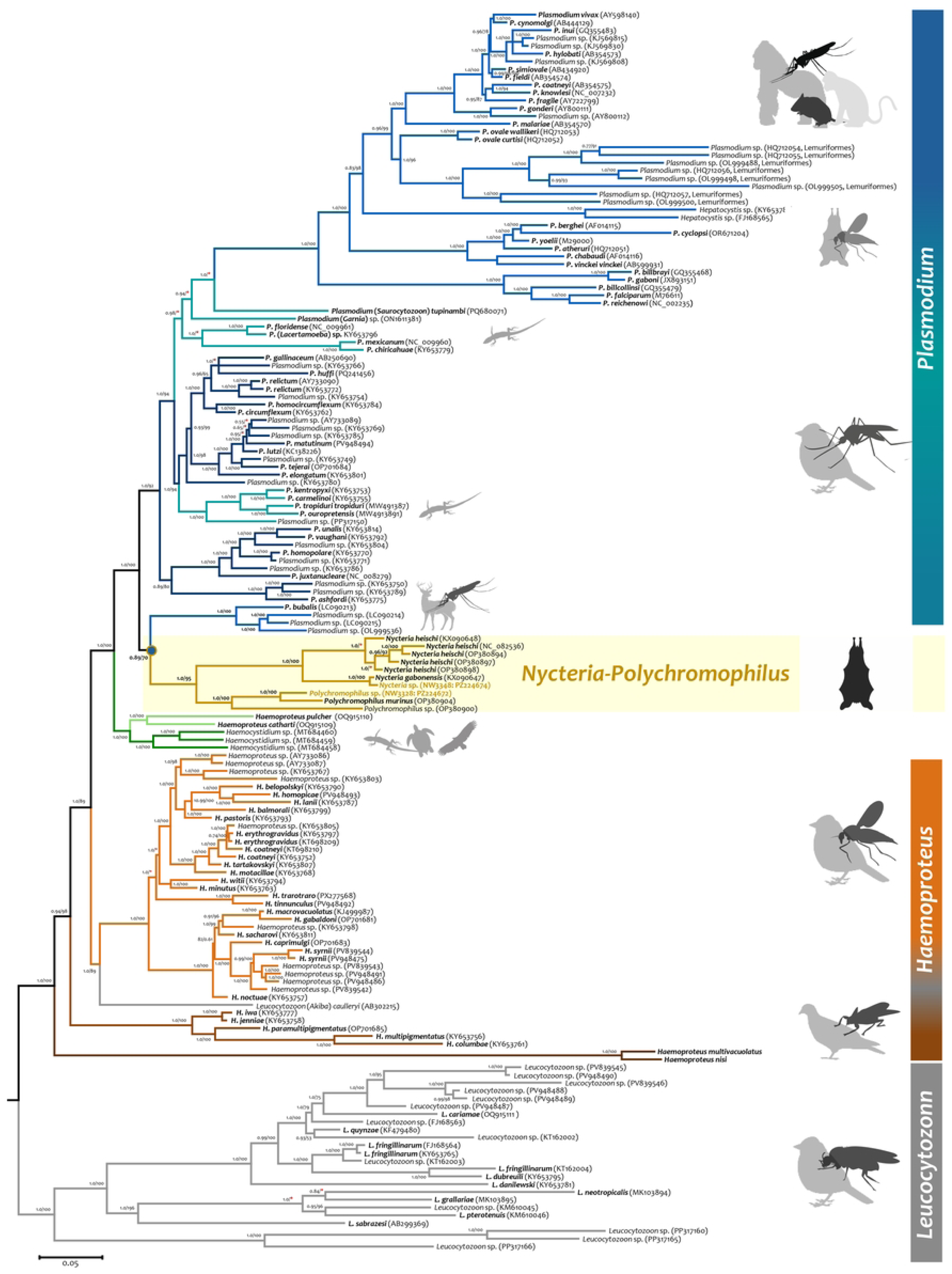
Bayesian phylogenetic hypothesis of haemosporidian species based on near-complete mitochondrial genomes (159 sequences and 5,052 bp excluding gaps). Node values represent posterior probabilities; bootstrap support from the ML tree is also reported as a percentage (see Materials and Methods). Branch colors indicate the genus, and red asterisks indicate inconsistencies between both phylogenetic methods (BI and ML). *Nycteria* and *Polychromophilus* parasites are highlighted with a light-yellow square, and the common ancestor of the Bovidae, *Nycteria*, and *Polychromophilus* parasites is indicated with a blue circle. GenBank accession numbers are given in parentheses.

The mitochondrial genome phylogeny recovered *Leucocytozoon*, *Haemoproteus*, and *Plasmodium* as polyphyletic (Fig 1), consistent with previous analyses of near-complete mitochondrial genomes [9,16,17,24]. In contrast, *Nycteria* and *Polychromophilus* formed a strongly supported monophyletic group (PP = 1.0; B = 95; Fig 1). This bat-parasite clade was sister to haemosporidians infecting the family Bovidae, but the node received lower support (PP = 0.89; B = 70). The Bovidae, *Nycteria*, and *Polychromophilus* parasites formed a sister clade with the *Plasmodium*-*Hepatocystis* species (PP = 1.0, B = 100). All these genera shared a common ancestor with the monophyletic clade comprising *Haemocystidium* spp., *Haemoproteus pulcher*, and *Haemoproteus catharti*. Within this topology, *Haemoproteus* (subgenus *Parahaemoproteus*) grouped with *Leucocytozoon (Akiba) caulleryi* (PP = 1.0; B = 89, Fig 1).

Furthermore, *Haemoproteus nisi* and *Haemoproteus multivacuolatus* were found to share a common ancestor (PP = 1.0, B = 100; Fig 1) with the clade comprising the genera *Plasmodium*, *Hepatocystis*, *Nycteria*, *Polychromophilus*, and *Haemocystidium*, as well as *L. caulleryi* and both subgenera of *Haemoproteus* (*Parahaemoproteus* and *Haemoproteus*). Interestingly, this mtDNA genome phylogeny corroborated the previously observed associations between specific parasite clades and vector groups at the family or subfamily level [9,25,26].

Phylogenies reconstructed from the three mitochondrial coding sequences (CDS: *cox3*, *cox1*, and *cytb* genes; 162 sequences, 3,270 bp excluding gaps) showed greater incongruence between BI and ML methods, with several poorly supported nodes (Fig 2). As with the near-complete mitochondrial phylogeny, only the BI topology is shown, with posterior probabilities (PP) and ML bootstrap values (B) reported for comparison. Many clades remained consistent with the near-complete mitochondrial phylogeny (Fig 1), including the non-monophyly of *Leucocytozoon*,

**Fig 2.**
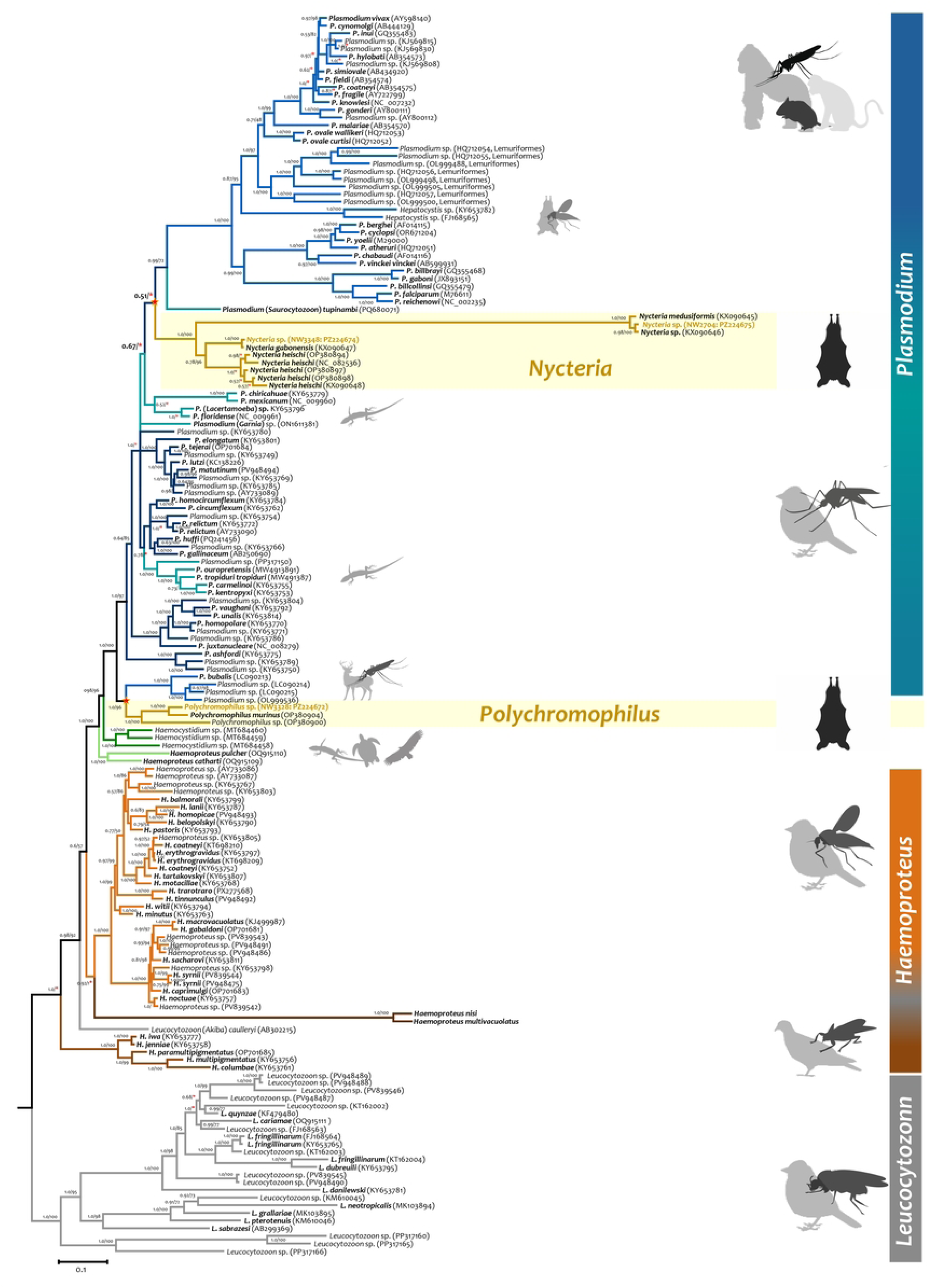
Bayesian phylogenetic hypothesis of haemosporidian species based only on the three mitochondrial coding sequences (CDS: *cox3*, *cox1*, and *cytb* genes; 162 sequences, 3,270 bp). The nodes’ values are posterior probabilities, and bootstrap support is reported as a percentage (see Materials and Methods). Branch colors indicate the genus, and red asterisks indicate inconsistencies between both phylogenetic methods (BI and ML). *Nycteria* and *Polychromophilus* parasites are highlighted with a light-yellow square. GenBank accession numbers are given in parentheses.

*Haemoproteus*, and *Plasmodium*. However, the genera *Nycteria* and *Polychromophilus* were not recovered as a monophyletic group. Specifically, *Nycteria* shared a common ancestor with primate/rodent *Plasmodium-Hepatocystis* species and one lizard *Plasmodium* parasite, albeit with low PP support (Fig 2), whereas *Polychromophilus* was placed as a sister group to the *Plasmodium* species infecting Bovidae with high support (PP=1.0, B=96; Fig 2). This discrepancy further illustrates the instability of *Nycteria*’s placement in reduced datasets.

Despite their particularly long branches, the three parasites within the genus *Nycteria* with rearranged mitochondrial genomes, *N. medusiformis* (KX090645), *Nycteria* sp. (KX090646) [21], and the new sequence *Nycteria* sp. (NW2704), formed a strongly supported monophyletic clade (PP = 1.0, B = 100; Fig 2). Internal relationships within *Nycteria* remained unresolved and showed substantial topological discordance between the BI and ML phylogenies (Fig 2). In contrast, the internal relationships within *Polychromophilus* were congruent across all analyses (BI and ML methods), with identical topologies and high node supports recovered from both the near-complete mtDNA genome and CDS-only datasets. Two additional topological discrepancies were observed between the mtDNA genome and CDS-only datasets: the phylogenetic placement of *Leucocytozoon (Akiba) caulleryi* and the *H. nisi/H. multivacuolatus* clade, as well as *Haemoproteus* subgenus *Haemoproteus* (Fig 2).

### Nuclear phylogeny based on nine loci and single-locus phylogenies

Phylogenetic inference on the concatenated alignment of nine nuclear loci (54 taxa, 6,543 bp excluding gaps) did not recover the monophyly of *Nycteria* and *Polychromophilus* (Fig 3). Instead, *Polychromophilus* groups with ungulate-infecting haemosporidians (PP = 1.0, B = 100; Fig 3). *Nycteria* shared a common ancestor with two major subclades: one comprising sauropsid *Plasmodium* taxa and the other containing mammalian *Plasmodium* and *Hepatocystis*. However, support for the sister relationship between sauropsid and mammalian *Plasmodium* lineages is weak (posterior probability, PP = 0.58).

**Fig 3.**
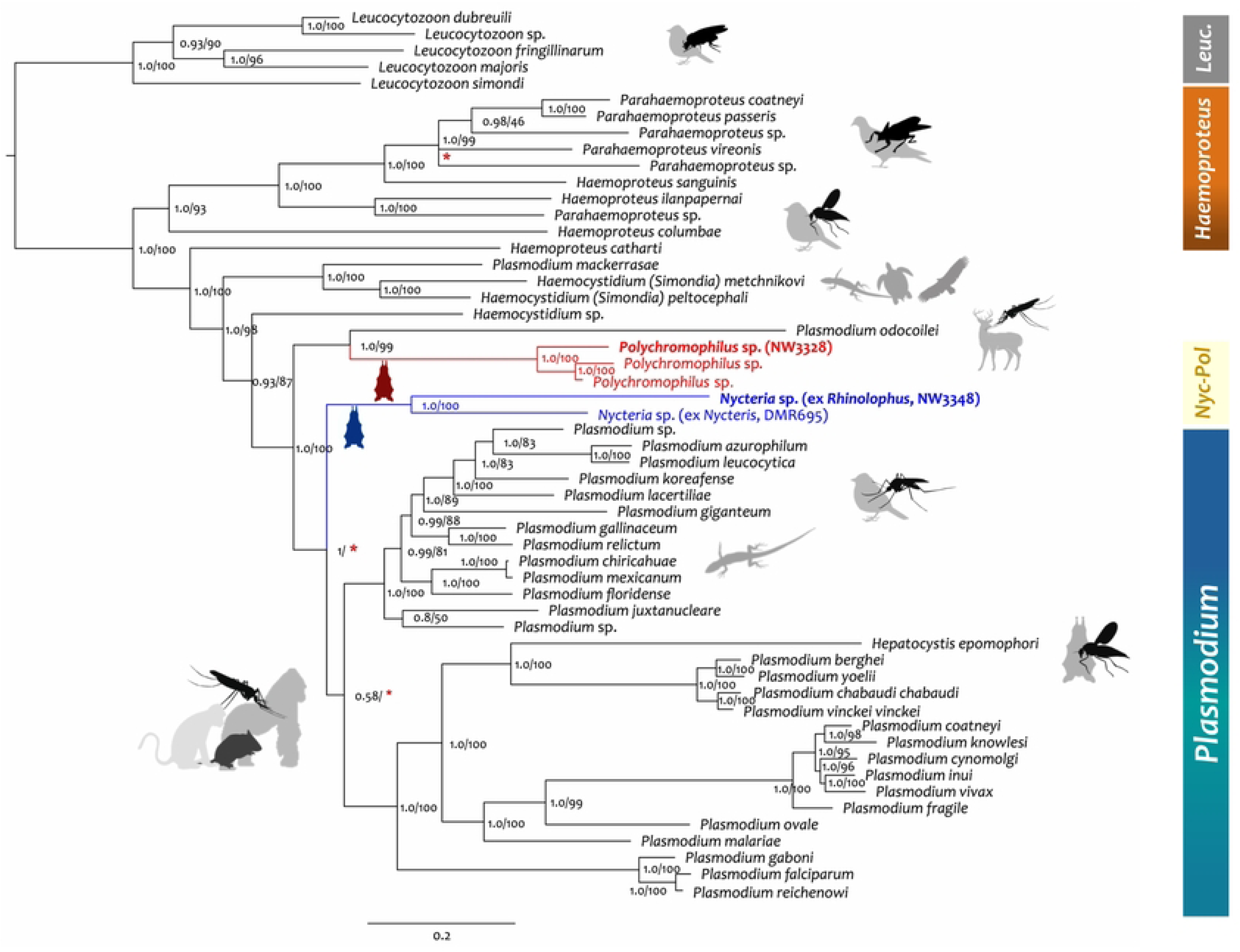
Bayesian phylogenetic hypothesis of haemosporidian species based on nine nuclear genes (54 sequences and 6,543 bp excluding gaps). Node values are posterior probabilities, and bootstrap support is reported as a percentage. Red asterisks indicate inconsistencies between both phylogenetic methods (BI and ML). GenBank accession numbers are given in the single gene trees in S1A-I Figure.

To evaluate locus-specific phylogenetic signal, each of the nine nuclear markers was analyzed individually using datasets with comparatively low levels of missing data (0.3-5.9%); individual gene trees are provided in S1 Fig. Across loci, topologies were highly variable and often exhibited extensive polytomies, particularly at deeper nodes. In several gene trees, relationships among major haemosporidian lineages remained unresolved, indicating limited phylogenetic signal at this taxonomic depth.

Marked discordance among loci was observed in the placement of *Nycteria* and *Polychromophilus*. The *Sec24* marker recovered *Nycteria, Polychromophilus*, and the ungulate-infecting *Plasmodium* taxon within a single clade (S1A Fig.); however, support for this grouping was low (PP = 0.77, B =54). In contrast, the *DrugT* locus recovered a markedly different topology, in which *Polychromophilus* was placed in a deep basal position, outside a clade containing *Simondia* and sauropsid *Plasmodium*, whereas *Nycteria* was placed basal to the mammalian *Plasmodium* clade (S1B Fig). The *POLD1* marker mirrored the concatenated topology with respect to the relative positions of *Polychromophilus* and *Nycteria* but exhibited extensive polytomies within the *Haemocystidium* and *Haemoproteus* (subgenus *Parahaemoproteus*) clades, limiting resolution among these taxa (S1C Fig). The remaining nuclear loci were largely dominated by polytomies and did not consistently resolve relationships among *Polychromophilus*, *Nycteria*, and other major haemosporidian lineages (S1D-I Fig).

### Timing the radiation of *Polychromophilus* and *Nycteria* genera using the mtDNA genome

Divergence times were estimated using the near-complete mtDNA alignment (159 parasite species), which provided higher phylogenetic resolution (Fig 1) compared to the three-coding-sequence (CDS) phylogeny (Fig 2). Molecular dating was performed using MCMCTree [27–29], a Bayesian approach, and a non-Bayesian framework implemented in RelTime [30]. In MCMCTree, divergence times were estimated under both autocorrelated and independent-rate relaxed-clock models. Because CorrTest [31] supported the autocorrelated model, only timetrees from the autocorrelated analyses are shown (Figs. 4 and 5). However, node age estimates and their 95% highest posterior density (HPD) intervals for major haemosporidian divergences under the three calibration scenarios and both rate models are summarized in Tables 1–3. Across methods, three calibration scenarios were evaluated, each including four to six calibration constraints defined by uniform distributions (see Materials and Methods).

**Fig 4.**
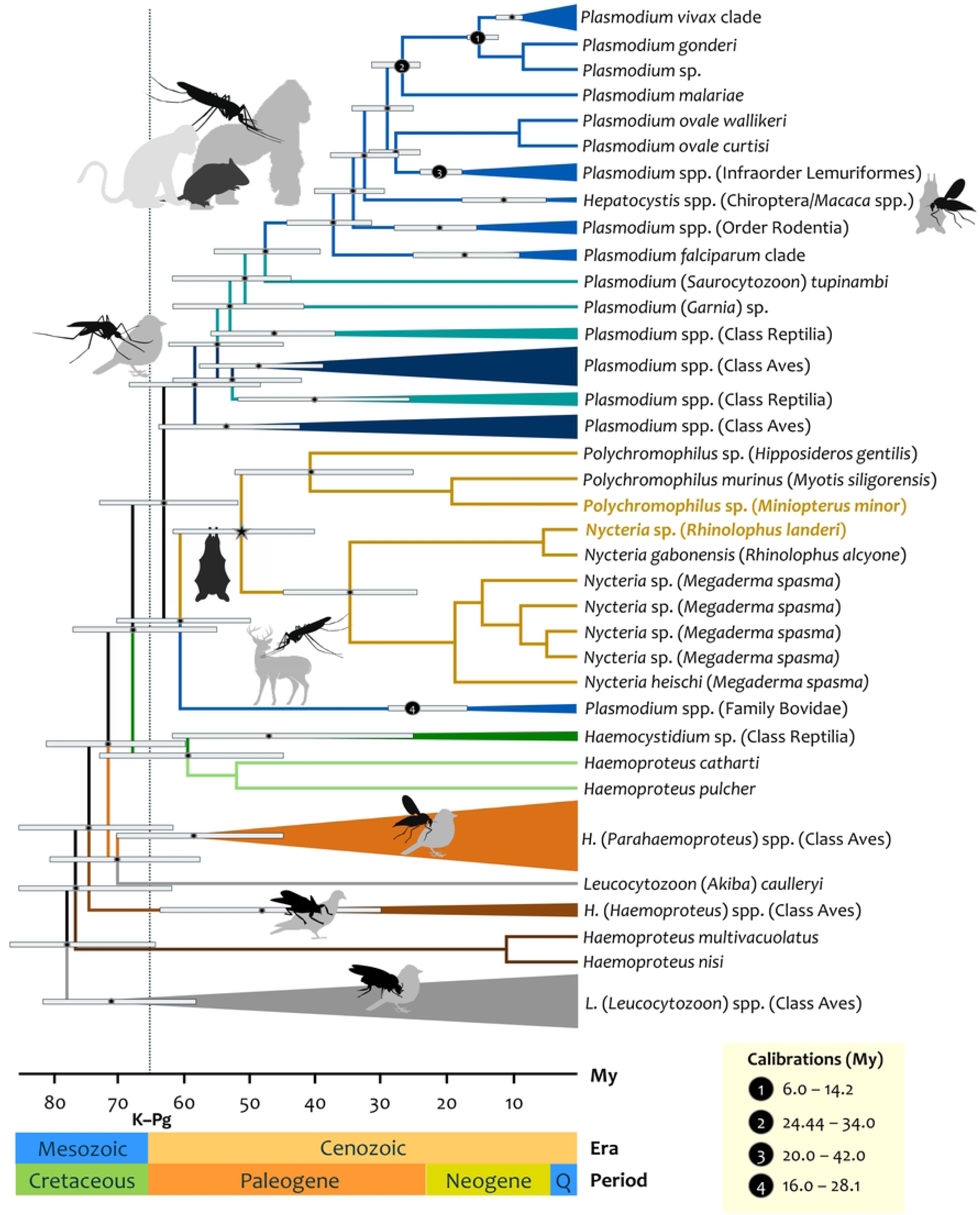
Divergence time estimates for the major haemosporidian clades under the first calibration scenario. Timetree was generated using MCMCTree method under the first calibration scenario with an autocorrelated rate model. Estimates are based on a mitochondrial genome alignment of 159 sequences (5,052 bp, excluding gaps and rearranged genomes). Branch colors denote different genera; bars at the nodes represent the 95% highest probability density (HPD). Novel sequences for *Nycteria* and *Polychromophilus* are highlighted in orange. Black circles indicate calibration constraints. All temporal estimates are expressed in millions of years (Ma).

**Fig 5.**
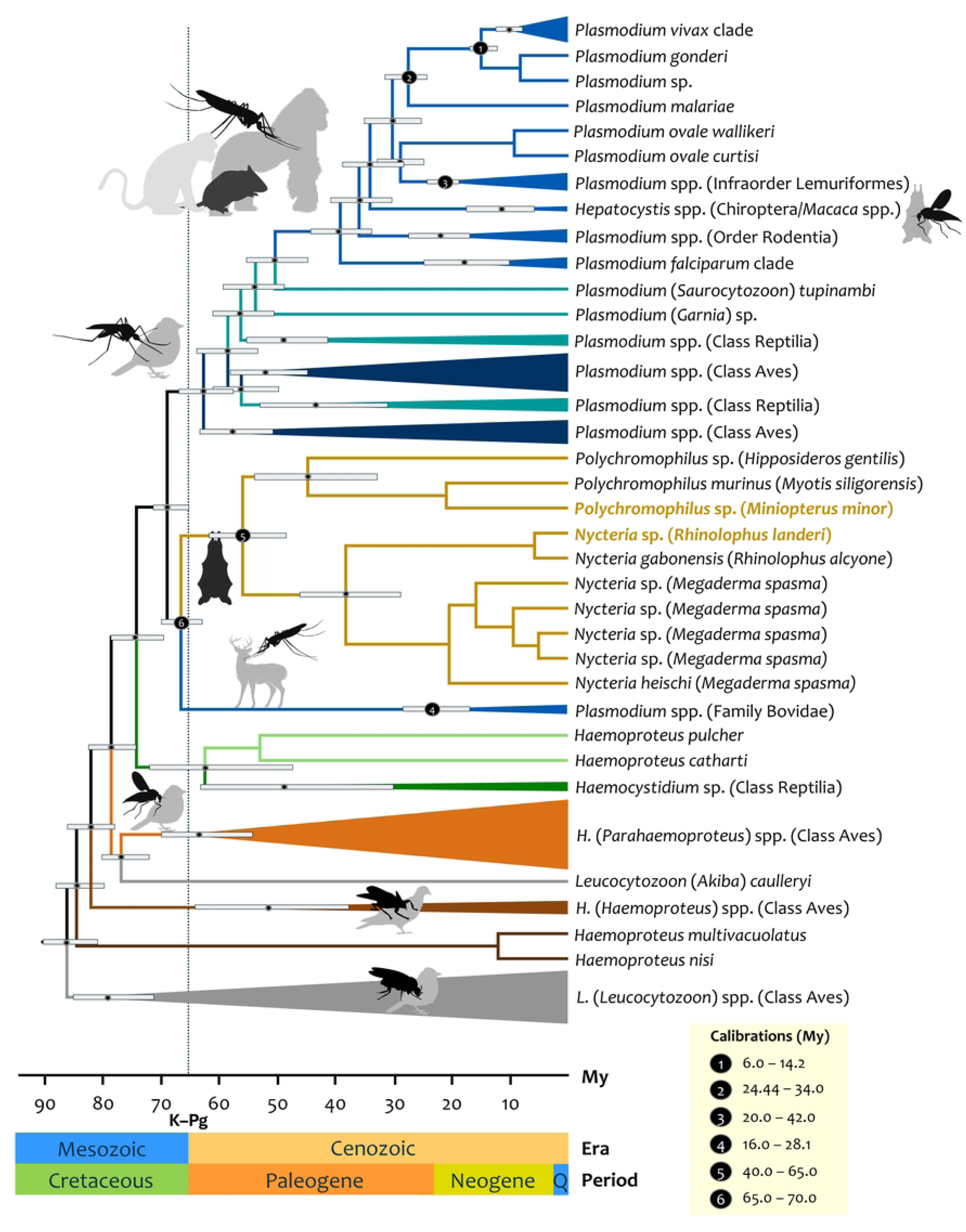
Divergence time estimates for major haemosporidian clades under the third calibration scenario. Timetree was generated using MCMCTree method under the third calibration scenario with an autocorrelated rate model. Estimates are based on a mitochondrial genome alignment of 159 sequences (5,052 bp, excluding gaps and rearranged genomes). Branch colors denote different genera; bars at the nodes represent the 95% highest probability density (HPD). Novel sequences for *Nycteria* and *Polychromophilus* are highlighted in orange. Black circles indicate calibration constraints. All temporal estimates are expressed in millions of years (Ma).

**Table 1.** Divergence time estimates for the major haemosporidian clades under the first calibration scenario.

| Divergence (Ma) | Node | Scenario 1 |  |  |  |  |  |
| --- | --- | --- | --- | --- | --- | --- | --- |
|  |  | Independent |  | Correlated |  | RelTime |  |
|  |  | Node age | 95% HPD | Node age | 95% HPD | Node age | 95% 95% CI |
| Origin of *Plasmodium* spp./*Plasmodium vivax* (human and Asian NHP) | 240 | **10.58** | 8.25-12.82 | **10.24** | 8.48-11.96 | **6.67** | 4.44-10.02 |
| Origin of *Plasmodium* spp. (Rodentia) | 261 | **20.65** | 14.03-27.42 | **21.20** | 15.78-27.50 | **13.90** | 6.63-29.13 |
| Origin of *Plasmodium* spp. (Lemuriformes) | 254 | **24.18** | 19.78-29.24 | **20.92** | 18.43-24.38 | **21.71** | 20.00-35.34 |
| Origin of *Laverania* | 266 | **21.31** | 13.84-28.20 | **17.09** | 9.59-24.76 | **12.73** | 5.74-28.24 |
| Origin of *Plasmodium* + *Hepatocystis* genera (primates/rodents/bats) | 233 | **39.07** | 32.35-60.58 | **37.03** | 31.09-43.73 | **31.29** | 24.44-58.44 |
| Origin of *Plasmodium*+ *Hepatocystis* spp. (Aves/Reptilia/Rodent/Primate) | 195 | **50.64** | 41.78-60.40 | **62.88** | 47.99-67.68 | **41.11** | 26.47-63.85 |
| Origin of *Polychromophilus* | 193 | **23.03** | 12.21-34.75 | **40.55** | 25.75-52.50 | **25.55** | 12.31-53.00 |
| Origin of *Nycteria* | 187 | **19.82** | 11.84-27.36 | **34.46** | 24.12-45.20 | **15.53** | 7.12-33.87 |
| Split between *Polychromophilus* and *Nycteria* | 186 | **36.53** | 24.27-48.75 | **51.04** | 40.08-61.67 | **34.47** | 19.76-60.13 |
| Origin of *Plasmodium* spp. (Family Bovidae) | 183 | **19.50** | 15.44-25.63 | **25.23** | 17.39-29.04 | **21.07** | 16.00-27.97 |
| Split between *Polychromophilus*/*Nycteria* and *Plasmodium* spp. (Family Bovidae) | 182 | **49.61** | 39.25-60.51 | **60.38** | 49.68-70.38 | **43.94** | 26.89-71.81 |
| Origin of *Plasmodium* + *Hepatocystis* + *Polychromophilus* + *Nycteria* | 181 | **53.89** | 44.33-64.09 | **62.88** | 52.16-72.98 | **46.82** | 29.63-74.00 |
| Origin of the genus *Haemocystidium* + (*H. pulcher*/*H. catharti*) | 177 | **25.27** | 14.05-40.60 | **59.23** | 44.92-73.02 | **45.67** | 26.89-77.58 |
| Split between *Plasmodium*/*Hepatocystis*/*Polychromophilus*/*Nycteria* and *Haemocystidium* | 176 | **57.58** | 47.59-68.68 | **67.62** | 56.31-77.93 | **53.61** | 32.73-86.80 |
| Origin of the genus *Haemoproteus* (*Parahaemoproteus*) | 145 | **25.98** | 19.44-34.16 | **58.39** | 45.83-69.89 | **40.64** | 22.37-73.84 |
| Split between *Haemoproteus* (*Parahaemoproteus*) and *L. caulleryi* | 144 | **57.70** | 45.96-70.68 | **69.91** | 58.34-80.51 | **60.00** | 34.59-86.80 |
| Origin of *Haemoproteus* (*Haemoproteus*) | 139 | **25.91** | 15.52-39.20 | **47.67** | 30.55-63.80 | **29.72** | 14.86-59.47 |
| Origin of *Leucocytozoon* (*Leucocytozoon*) | N/A | **68.4** | 54.81-82.46 | **70.95** | 57.65-82.87 | **N/A** | N/A |
| Origin of Haemosporida | N/A | **73.8** | 61.47-86.94 | **77.66** | 65.17-87.75 | **N/A** | N/A |

Values were generated using MCMCTree (autocorrelated and independent rates models) and RelTime under the first calibration scenario. Point estimates are provided in millions of years (Ma) with associated 95% highest probability density (HPD) for MCMCTree and confidence intervals (CI) for RelTime. Calibration constraints: (1) The Papio/Macaca parasite divergence: 6.0–14.2 Ma; (2) the Human/Macaca split: 24.44–34.0 Ma; (3) the origin of Lemuridae–Indriidae clade: 20.0–42.0 Ma; and (4) the Bovinae–Antilopinae divergence: 16.0–28.1 Ma. N/A, not applicable.

**Table 2.**
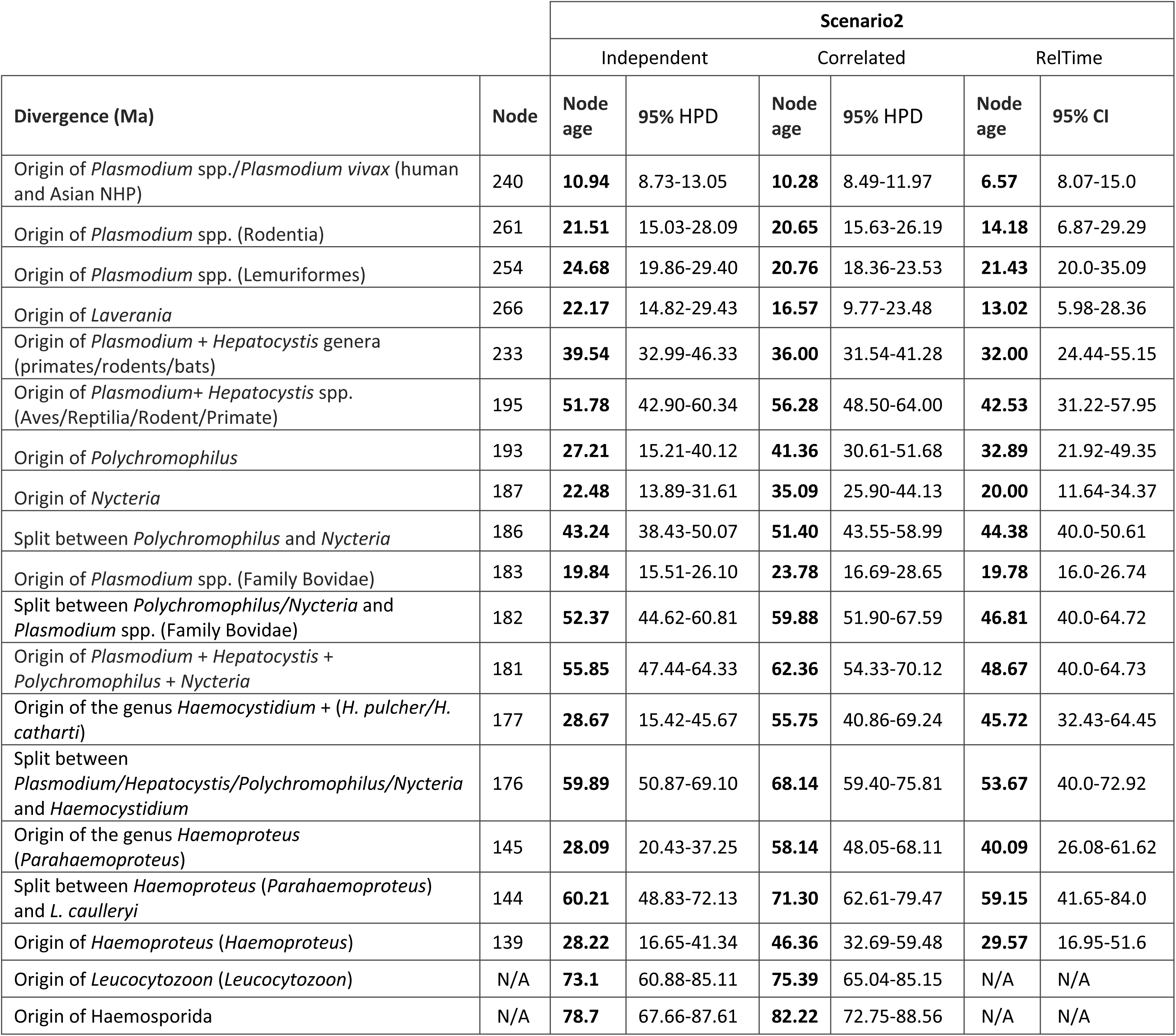
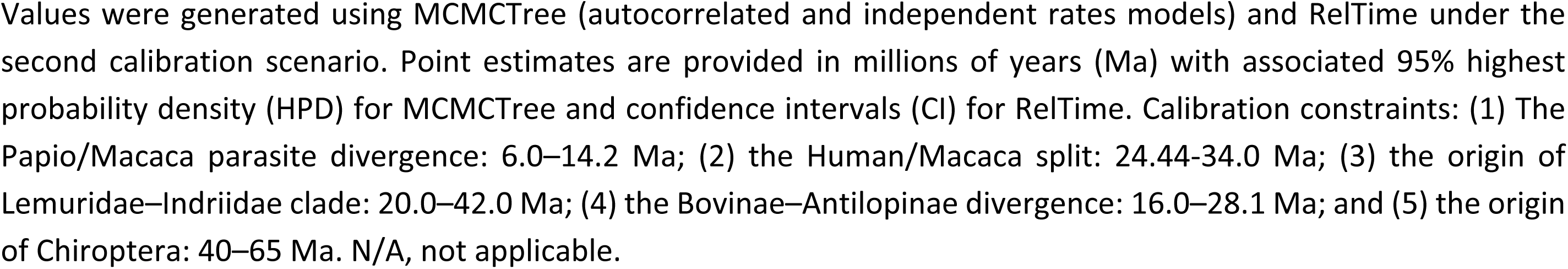
Divergence time estimates for the major haemosporidian clades under the second calibration scenario.

| Divergence (Ma) | Node | Scenario2 |  |  |  |  |  |
| --- | --- | --- | --- | --- | --- | --- | --- |
|  |  | Independent |  | Correlated |  | RelTime |  |
|  |  | Node age | 95% HPD | Node age | 95% HPD | Node age | 95% CI |
| Origin of *Plasmodium* spp./*Plasmodium vivax* (human and Asian NHP) | 240 | **10.94** | 8.73-13.05 | **10.28** | 8.49-11.97 | **6.57** | 8.07-15.0 |
| Origin of *Plasmodium* spp. (Rodentia) | 261 | **21.51** | 15.03-28.09 | **20.65** | 15.63-26.19 | **14.18** | 6.87-29.29 |
| Origin of *Plasmodium* spp. (Lemuriformes) | 254 | **24.68** | 19.86-29.40 | **20.76** | 18.36-23.53 | **21.43** | 20.0-35.09 |
| Origin of *Laverania* | 266 | **22.17** | 14.82-29.43 | **16.57** | 9.77-23.48 | **13.02** | 5.98-28.36 |
| Origin of *Plasmodium* + *Hepatocystis* genera (primates/rodents/bats) | 233 | **39.54** | 32.99-46.33 | **36.00** | 31.54-41.28 | **32.00** | 24.44-55.15 |
| Origin of *Plasmodium*+ *Hepatocystis* spp. (Aves/Reptilia/Rodent/Primate) | 195 | **51.78** | 42.90-60.34 | **56.28** | 48.50-64.00 | **42.53** | 31.22-57.95 |
| Origin of *Polychromophilus* | 193 | **27.21** | 15.21-40.12 | **41.36** | 30.61-51.68 | **32.89** | 21.92-49.35 |
| Origin of *Nycteria* | 187 | **22.48** | 13.89-31.61 | **35.09** | 25.90-44.13 | **20.00** | 11.64-34.37 |
| Split between *Polychromophilus* and *Nycteria* | 186 | **43.24** | 38.43-50.07 | **51.40** | 43.55-58.99 | **44.38** | 40.0-50.61 |
| Origin of *Plasmodium* spp. (Family Bovidae) | 183 | **19.84** | 15.51-26.10 | **23.78** | 16.69-28.65 | **19.78** | 16.0-26.74 |
| Split between *Polychromophilus*/*Nycteria* and *Plasmodium* spp. (Family Bovidae) | 182 | **52.37** | 44.62-60.81 | **59.88** | 51.90-67.59 | **46.81** | 40.0-64.72 |
| Origin of *Plasmodium* + *Hepatocystis* + *Polychromophilus* + *Nycteria* | 181 | **55.85** | 47.44-64.33 | **62.36** | 54.33-70.12 | **48.67** | 40.0-64.73 |
| Origin of the genus *Haemocystidium* + (*H. pulcher*/*H. catharti*) | 177 | **28.67** | 15.42-45.67 | **55.75** | 40.86-69.24 | **45.72** | 32.43-64.45 |
| Split between *Plasmodium*/*Hepatocystis*/*Polychromophilus*/*Nycteria* and *Haemocystidium* | 176 | **59.89** | 50.87-69.10 | **68.14** | 59.40-75.81 | **53.67** | 40.0-72.92 |
| Origin of the genus *Haemoproteus* (*Parahaemoproteus*) | 145 | **28.09** | 20.43-37.25 | **58.14** | 48.05-68.11 | **40.09** | 26.08-61.62 |
| Split between *Haemoproteus* (*Parahaemoproteus*) and *L. caulleryi* | 144 | **60.21** | 48.83-72.13 | **71.30** | 62.61-79.47 | **59.15** | 41.65-84.0 |
| Origin of *Haemoproteus* (*Haemoproteus*) | 139 | **28.22** | 16.65-41.34 | **46.36** | 32.69-59.48 | **29.57** | 16.95-51.6 |
| Origin of *Leucocytozoon* (*Leucocytozoon*) | N/A | **73.1** | 60.88-85.11 | **75.39** | 65.04-85.15 | N/A | N/A |
| Origin of Haemosporida | N/A | **78.7** | 67.66-87.61 | **82.22** | 72.75-88.56 | N/A | N/A |

Values were generated using MCMCTree (autocorrelated and independent rates models) and RelTime under the second calibration scenario. Point estimates are provided in millions of years (Ma) with associated 95% highest probability density (HPD) for MCMCTree and confidence intervals (CI) for RelTime. Calibration constraints: (1) The Papio/Macaca parasite divergence: 6.0–14.2 Ma; (2) the Human/Macaca split: 24.44–34.0 Ma; (3) the origin of Lemuridae–Indriidae clade: 20.0–42.0 Ma; (4) the Bovinae–Antilopinae divergence: 16.0–28.1 Ma; and (5) the origin of Chiroptera: 40–65 Ma. N/A, not applicable.

**Table 3.**
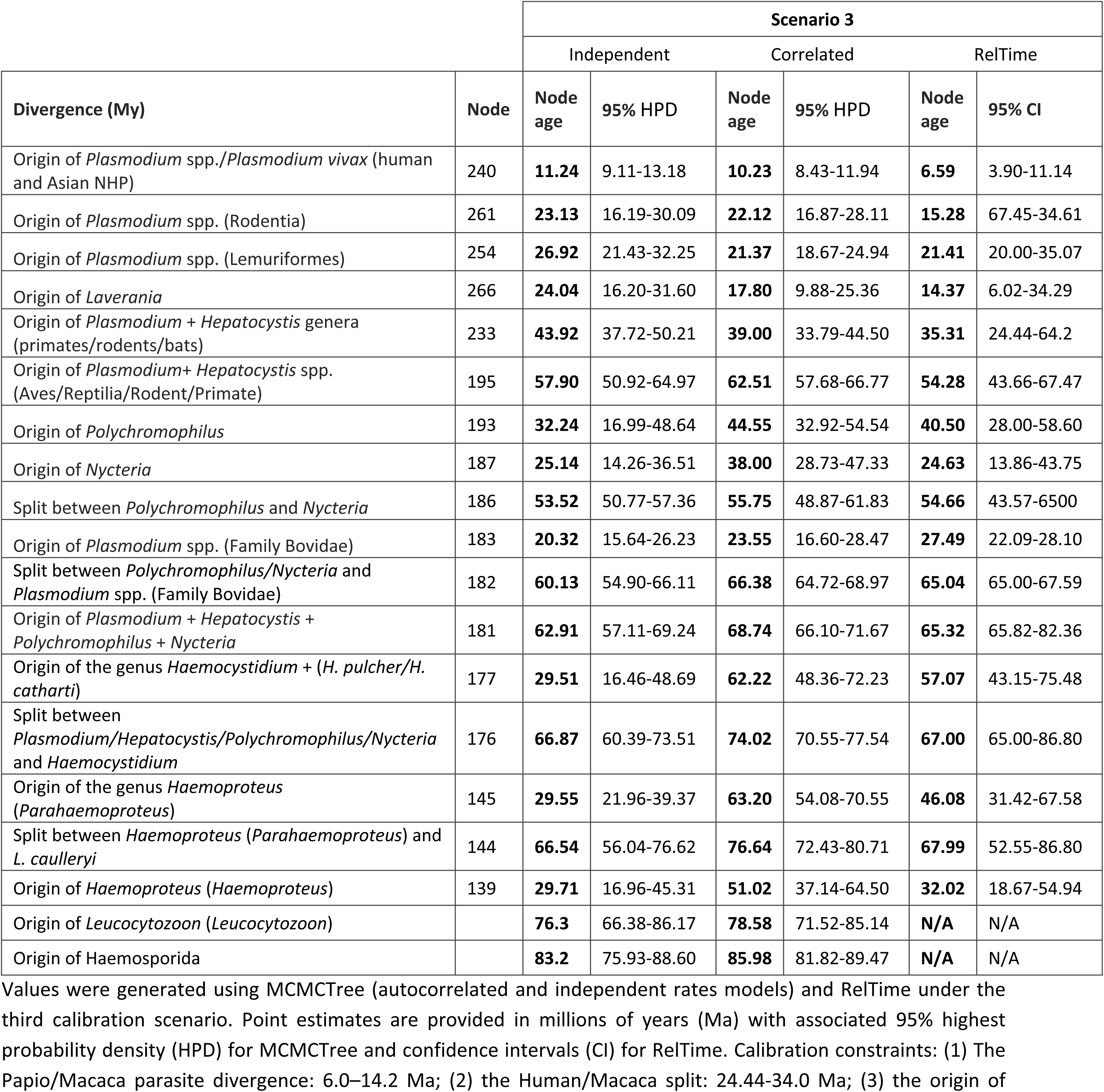
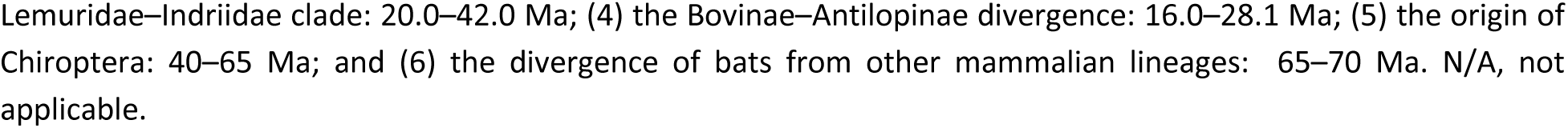
Divergence time estimates for the major haemosporidian clades under the third calibration scenario.

| Divergence (My) | Node | Scenario 3 |  |  |  |  |  |
| --- | --- | --- | --- | --- | --- | --- | --- |
|  |  | Independent |  | Correlated |  | RelTime |  |
|  |  | Node age | 95% HPD | Node age | 95% HPD | Node age | 95% CI |
| Origin of *Plasmodium* spp./*Plasmodium vivax* (human and Asian NHP) | 240 | **11.24** | 9.11-13.18 | **10.23** | 8.43-11.94 | **6.59** | 3.90-11.14 |
| Origin of *Plasmodium* spp. (Rodentia) | 261 | **23.13** | 16.19-30.09 | **22.12** | 16.87-28.11 | **15.28** | 67.45-34.61 |
| Origin of *Plasmodium* spp. (Lemuriformes) | 254 | **26.92** | 21.43-32.25 | **21.37** | 18.67-24.94 | **21.41** | 20.00-35.07 |
| Origin of *Laverania* | 266 | **24.04** | 16.20-31.60 | **17.80** | 9.88-25.36 | **14.37** | 6.02-34.29 |
| Origin of *Plasmodium* + *Hepatocystis* genera (primates/rodents/bats) | 233 | **43.92** | 37.72-50.21 | **39.00** | 33.79-44.50 | **35.31** | 24.44-64.2 |
| Origin of *Plasmodium*+ *Hepatocystis* spp. (Aves/Reptilia/Rodent/Primate) | 195 | **57.90** | 50.92-64.97 | **62.51** | 57.68-66.77 | **54.28** | 43.66-67.47 |
| Origin of *Polychromophilus* | 193 | **32.24** | 16.99-48.64 | **44.55** | 32.92-54.54 | **40.50** | 28.00-58.60 |
| Origin of *Nycteria* | 187 | **25.14** | 14.26-36.51 | **38.00** | 28.73-47.33 | **24.63** | 13.86-43.75 |
| Split between *Polychromophilus* and *Nycteria* | 186 | **53.52** | 50.77-57.36 | **55.75** | 48.87-61.83 | **54.66** | 43.57-6500 |
| Origin of *Plasmodium* spp. (Family Bovidae) | 183 | **20.32** | 15.64-26.23 | **23.55** | 16.60-28.47 | **27.49** | 22.09-28.10 |
| Split between *Polychromophilus*/*Nycteria* and *Plasmodium* spp. (Family Bovidae) | 182 | **60.13** | 54.90-66.11 | **66.38** | 64.72-68.97 | **65.04** | 65.00-67.59 |
| Origin of *Plasmodium* + *Hepatocystis* + *Polychromophilus* + *Nycteria* | 181 | **62.91** | 57.11-69.24 | **68.74** | 66.10-71.67 | **65.32** | 65.82-82.36 |
| Origin of the genus *Haemocystidium* + (*H. pulcher*/*H. catharti*) | 177 | **29.51** | 16.46-48.69 | **62.22** | 48.36-72.23 | **57.07** | 43.15-75.48 |
| Split between *Plasmodium*/*Hepatocystis*/*Polychromophilus*/*Nycteria* and *Haemocystidium* | 176 | **66.87** | 60.39-73.51 | **74.02** | 70.55-77.54 | **67.00** | 65.00-86.80 |
| Origin of the genus *Haemoproteus* (*Parahaemoproteus*) | 145 | **29.55** | 21.96-39.37 | **63.20** | 54.08-70.55 | **46.08** | 31.42-67.58 |
| Split between *Haemoproteus* (*Parahaemoproteus*) and *L. caulleryi* | 144 | **66.54** | 56.04-76.62 | **76.64** | 72.43-80.71 | **67.99** | 52.55-86.80 |
| Origin of *Haemoproteus* (*Haemoproteus*) | 139 | **29.71** | 16.96-45.31 | **51.02** | 37.14-64.50 | **32.02** | 18.67-54.94 |
| Origin of *Leucocytozoon* (*Leucocytozoon*) |  | **76.3** | 66.38-86.17 | **78.58** | 71.52-85.14 | **N/A** | N/A |
| Origin of Haemosporida |  | **83.2** | 75.93-88.60 | **85.98** | 81.82-89.47 | **N/A** | N/A |

Values were generated using MCMCTree (autocorrelated and independent rates models) and RelTime under the third calibration scenario. Point estimates are provided in millions of years (Ma) with associated 95% highest probability density (HPD) for MCMCTree and confidence intervals (CI) for RelTime. Calibration constraints: (1) The Papio/Macaca parasite divergence: 6.0–14.2 Ma; (2) the Human/Macaca split: 24.44–34.0 Ma; (3) the origin of
Lemuridae–Indriidae clade: 20.0–42.0 Ma; (4) the Bovinae–Antilopinae divergence: 16.0–28.1 Ma; (5) the origin of Chiroptera: 40–65 Ma; and (6) the divergence of bats from other mammalian lineages: 65–70 Ma. N/A, not applicable.

The first scenario utilized four calibration constraints previously tested by [9,16,17]. Although divergence estimates for the origins of *Nycteria* and *Polychromophilus* varied across the methods and rate models, their HPD intervals exhibited significant overlap (Table 1, Fig 4). Notably, the results from the independent rates model in MCMCTree were closely aligned with those generated by RelTime (S2 Fig). In contrast, the autocorrelated model produced significantly older estimates (S2 Fig); specifically, the origin of *Polychromophilus* was dated to 40.55 Ma (HDP: 25.75–52.50) and that of *Nycteria* to 34.46 Ma (HDP: 24.12–45.20). Interestingly, their HDP intervals overlapped with the established evolutionary origins of their chiropteran hosts [32,33]. Based on these results, two additional calibration scenarios were explored to further test the robustness of the findings.

Divergence-time estimates for the major haemosporidian clades under the second scenario (Table 2), which incorporated a 40–65 Ma uniform prior for the origin of chiropteran parasites, were highly consistent with those obtained under the first scenario (Table 1, S2 Fig). The estimated split between *Nycteria* and *Polychromophilus* was 51.40 Ma (HPD: 43.55–58.99 Ma), nearly identical to the estimate under scenario 1 (51.04 Ma; HPD: 40.08–61.67 Ma). Likewise, the divergence between the *Nycteria–Polychromophilus* clade and *Plasmodium* species infecting Bovidae was highly similar across both scenarios, with estimates of 59.88 Ma (HPD: 51.90–67.59 Ma) and 60.38 Ma (HPD: 49.68–70.38 Ma), respectively, overlapping with proposed times for early bat diversification [32,33].

The addition of a sixth calibration constraint (65–70 Ma) in the third scenario yielded divergence-time estimates that remained broadly consistent with those obtained under the first two scenarios (Table 3, Fig 5). To assess the influence of these calibrations on time estimates, RelTime was used with and without calibration constraints, following the protocol of Battistuzzi et al. [34]. Across all scenarios, comparisons of calibrated and uncalibrated estimates yielded high correlation coefficients, with the third scenario showing a slightly better fit (R^2^ = 0.99 vs. 0.98, S3 Fig). These results indicate that the six calibration constraints, including the newly proposed ones, are internally consistent. The HPD intervals obtained under the third scenario (Fig 5) were also narrower than those produced under scenarios 1 and 2. Collectively, the three calibration scenarios yielded broadly consistent divergence-time estimates, with the third scenario producing the narrowest HPD intervals (Fig 5, Tables 1–3).

Branch-specific relative evolutionary rates were estimated for the near-complete mtDNA with RelTime (Fig 6). These analyses show heterogeneity in rates across haemosporidian lineages. *Nycteria* lineages exhibited elevated substitution rates relative to other mammalian haemosporidians, consistent with their long branches in the phylogeny. In contrast, *Polychromophilus* showed a more moderate rate of variation, with branch lengths comparable to those of other mammal-infecting taxa. Other clades with accelerated substitution rates included: (1) the rodent-bat-primate *Plasmodium* lineage, most notably in *Plasmodium cyclopsi* (OR671204, sampled from the bat host *Doryrhina cyclops*); (2) *Haemoproteus* (*Parahaemoproteus*) spp. infecting non-passerine hosts, such as *Haemoproteus gabaldoni* (Anatidae) and *Haemoproteus syrnii* (Strigiformes); and (3) *Haemoproteus* (*Haemoproteus*) spp. of non-passerines, including *Haemoproteus columbae* and *Haemoproteus multipigmentatus* (Columbiformes).

**Fig 6.**
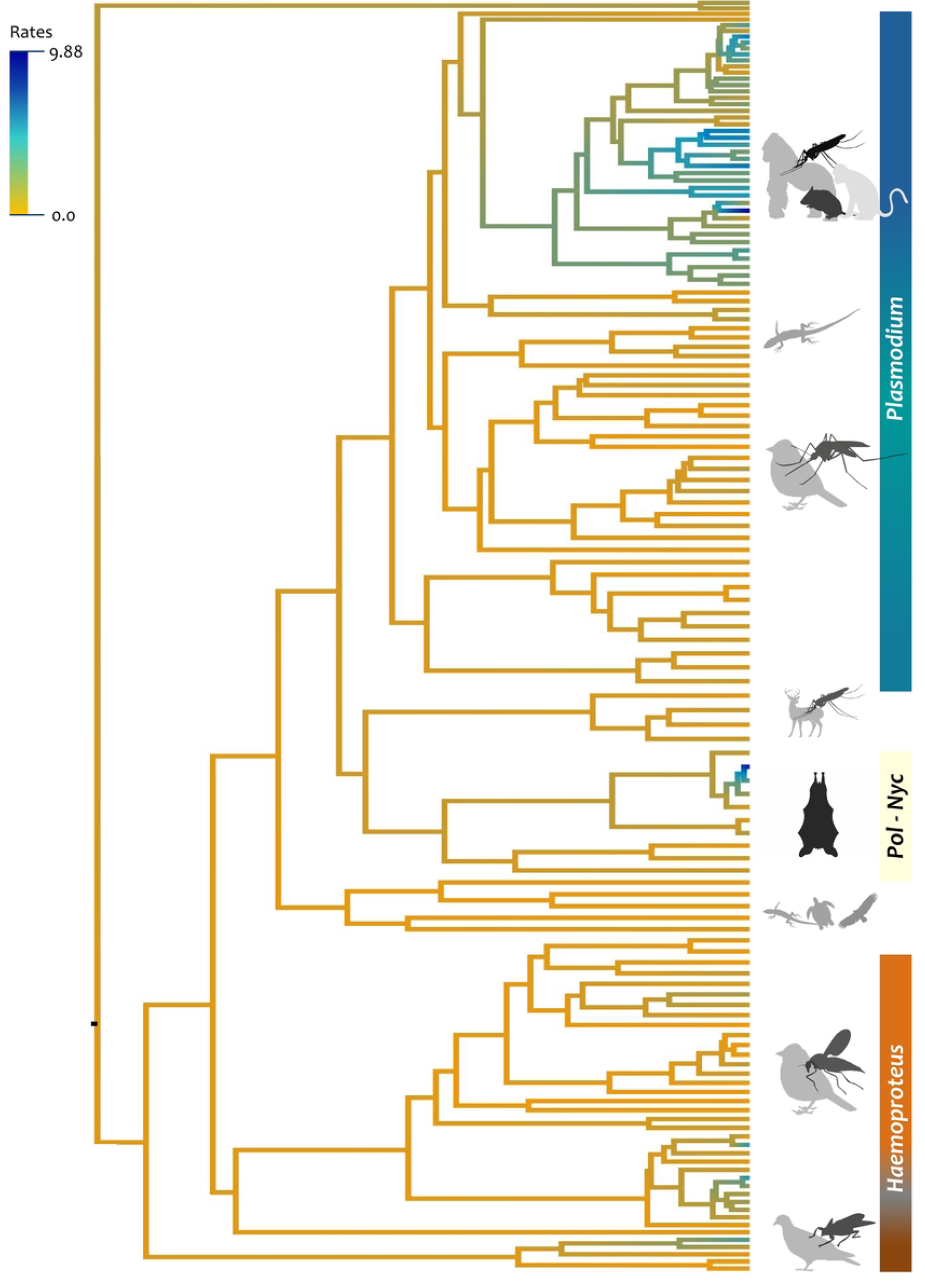
Heterogeneity in evolutionary rates across haemosporidian mitochondrial genomes. Branch colors represent relative substitution rates, ranging from high (blue) to low (light orange), normalized to the root rate (set to unity). Rates were estimated using the RelTime method in MEGA without calibration constraints. Taxonomic genera are color-coded in the legend on the right.

## Discussion

### Limited resolution of deep haemosporidian relationships using partial nuclear sequences

Although there are multiple phylogenetic inferences for Haemosporida, taxon sampling and data quality hamper their comparison (as revised in [1]). In our analyses, the near-complete mtDNA dataset recovered a stable and strongly supported *Nycteria* + *Polychromophilus* clade, whereas the CDS-only dataset showed greater method-dependent instability and failed to recover their monophyly. This discrepancy is better interpreted as a difference in the recovered phylogenetic signal, as the fragmented rRNA and other non-coding regions are absent from the coding-sequence alignment. In our case, the contrast is further complicated by the inclusion of the rearranged, long-branched *Nycteria* genomes in the CDS-only alignment, which likely contributes to shifts in *Nycteria*’s placement in the phylogeny. We therefore favor the interpretation that the near-complete mtDNA genome recovers phylogenetic signal more effectively, whereas the CDS-only topology reflects reduced character sampling and lineage-specific rate heterogeneity [9], particularly within *Nycteria*.

The reduced nine-locus nuclear dataset provides only limited resolution of deep relationships among major haemosporidian lineages. Although the concatenated nuclear analysis (6,543 bp excluding gaps) broadly recovered the major clades, relationships among *Nycteria*, *Polychromophilus*, ungulate *Plasmodium*, and the avian and mammalian *Plasmodium* lineages varied across loci and were often weakly supported. Many single-gene trees also exhibited extensive polytomies and low support at deeper nodes, indicating that this dataset contains insufficient phylogenetic signal at this taxonomic depth. The combination of low support, locus-specific topological conflict, and frequent polytomies is consistent with limited signal and underlying gene-tree heterogeneity [35]. Although concatenation can improve apparent resolution, it does not remove conflict among loci, and even with this number of sites, several critical nodes remain unresolved.

Re-examination of the alignment used by Galen et al. [10] revealed extensive missing data and limited overlap among comparable sites across taxa, resulting from uneven locus representation across haemosporidian lineages. In this context, sparse data among taxa can affect estimates of branch lengths and rate-heterogeneity parameters, potentially inflating support for relationships that are only weakly informed by shared characters [35–37]. To minimize these effects, we used carefully curated alignments with reduced missing data and a more comparable set of sites/loci across taxa. Importantly, all gene alignments in the present study included both *Nycteria* and *Polychromophilus* for every locus, ensuring that these genera were consistently represented across all markers. Although the concatenated dataset still contained 26.6% missing data overall due to the absence of sequences in other haemosporidian taxa, the sampling for our target genera was complete. Taken together, these results suggest that the limited informative signal in the current nuclear dataset, due to an uneven sampling of loci across the order, is the primary factor underlying inconsistent phylogenies between nuclear and mtDNA data.

Mitochondrial genomes have proven phylogenetically informative in haemosporidian systematics because they are compact and retain substantial signal across the order [9,16,17,24]. In our analysis, the near-complete mitochondrial dataset recovered the most stable and strongly supported topology. However, this result should not be treated as a definitive species-tree hypothesis. Mitochondrial DNA represents a single locus, and agreement with mtDNA + apicoplast datasets does not exclude deeper discordance across the nuclear genome [17]. A more definitive resolution of deep relationships within Haemosporida will require broader phylogenomic sampling across all major lineages, enabling explicit species-tree analyses and direct evaluation of gene-tree heterogeneity. Unfortunately, such data remains lacking, and current multilocus datasets do not provide a consistent representation of all clades. Our findings, therefore, refine the mitochondrial framework for understanding parasite diversification and propose that bats host a monophyletic and evolutionarily distinct haemosporidian radiation.

### History of phylogenetic placements of *Nycteria* and *Polychromophilus*

The instability observed in our nuclear analyses mirrors a broader pattern in previous phylogenetic studies of *Nycteria* and *Polychromophilus*, in which their placement has varied with taxon sampling, marker choice, and the analytical framework. The phylogenetic position of *Polychromophilus* has remained particularly unstable since the first molecular analyses. Early studies based largely on mitochondrial *cytb* gene placed *Polychromophilus* with sauropsid *Plasmodium,* suggesting an origin from bird- or reptile-infecting haemosporidians. However, its precise position remained unresolved with *cytb* data alone [38–40]. Consistent with this interpretation, Witsenburg et al. [41] proposed that the ancestor of *Polychromophilus* was likely a bird- or reptile-infecting *Plasmodium* lineage that subsequently switched to bats, accompanied by a shift in dipteran vector, using expanded taxon sampling and multiple molecular markers from the three parasite genomes.

However, alternative topologies have also been recovered. The multi-gene phylogeny by Schaer et al. [13] instead placed *Polychromophilus* as the basal lineage within a clade of exclusively mammal-infecting haemosporidians, highlighting the sensitivity of its placement to taxon sampling and analytical approach. Martinsen et al. [14] recovered white-tailed deer malaria parasites as sister taxon to *Polychromophilus*, whereas Templeton et al. [15] reinforced the divergence of ungulate malaria parasites as a monophyletic group but failed to fully resolve their phylogenetic relationship to *Polychromophilus*.

Similarly, analyses based on 21 nuclear loci, with limited informative sites due to the absence of common sampled loci across all taxa, recovered the deer parasite *Plasmodium odocoilei* as sister to *Polychromophilus*, with this lineage positioned closer to avian parasites than to other mammalian *Plasmodium* clades [10]. The phylogenetic placement of *Nycteria* has been equally controversial. The earliest sequence attributed to this lineage, obtained from a parasite infecting *Megaderma* bats, was initially recovered within the sauropsid *Plasmodium* clade, although the authors refrained from assigning it to a specific genus [38]. The first dedicated molecular study of *Nycteria,* based on a multilocus dataset, supported the genus as a valid and monophyletic lineage, using newly generated sequences from African bats that grouped with the previously reported *Megaderma*-derived sequence [13]. In that analysis, *Nycteria* formed a highly distinct lineage sister to the mammalian *Plasmodium*/*Hepatocystis* clade, highlighting its deep evolutionary divergence within the Haemosporida.

Subsequent analyses incorporating additional *Nycteria* taxa yielded conflicting placements. Schaer et al. [19] reported difficulties in amplifying mitochondrial markers for some *Nycteria* lineages and detected unusually high substitution rates in the cytochrome oxidase I (*cox1*) gene, suggesting substantial modifications to the mitochondrial genome. In that study, Bayesian analyses placed *Nycteria* and *Polychromophilus* within the sauropsid *Plasmodium* parasites, whereas maximum-likelihood analyses weakly recovered *Nycteria* as the sister to mammalian *Plasmodium*. Further work showed that some *Nycteria* species possess rearranged mitochondrial genomes, which explains earlier amplification failures and contributes to phylogenetic instability [21]. Similarly, Galen et al. [10] recovered *Nycteria* either as sister to the primary mammalian *Plasmodium*/*Hepatocystis* clade or as sister to sauropsid *Plasmodium*, depending on the analytical framework and model applied.

In contrast to these early conflicting topologies, our mitochondrial genome analyses recover *Nycteria* and *Polychromophilus* as a strongly supported monophyletic lineage that is sister to the haemosporidians infecting Bovidae. This result suggests that the previously observed associations with *Plasmodium* might be artifacts of reduced character sampling or long-branch attraction, particularly given the significantly accelerated evolutionary rates observed within the *Nycteria* lineage (Fig 6).

### Implications for malaria parasite evolution

Our divergence-time estimates provide a robust temporal framework for understanding these evolutionary transitions. The origin of the *Nycteria–Polychromophilus* clade was estimated at approximately 51.04 Ma (HPD: 40.08-61.67; correlated Scenario 1), a period that coincides with the early radiation of bats during the Paleocene–Eocene transition [32,33]. This temporal alignment suggests that the association between haemosporidians and chiropterans is ancient and likely stems from an early host-switching event from a common ancestor shared with parasites infecting ungulates. Furthermore, the absence of erythrocytic schizogony—a defining biological feature shared by both *Nycteria* and *Polychromophilus*—may represent an ancestral trait retained from this early radiation or a convergent adaptation to the specialized physiological environment of bat hosts [13,18]. This finding does not imply a single origin of Haemosporida parasitism in bats, as *Plasmodium* and *Hepatocystis* also infect bats; rather, it suggests that chiropterans have been repeatedly colonized by evolutionarily distinct lineages.

Despite their shared absence of erythrocytic schizogony [e.g., 18,42], the two genera exhibit striking biological divergence: *Polychromophilus* undergoes tissue development within the endothelial system, whereas *Nycteria* asexually develops in the liver [18,42,43]. *Polychromophilus* is transmitted by nycteribiid bat flies, obligate hematophagous ectoparasites of bats; however, the vector for *Nycteria* parasites remains unresolved [44–47]. Given that all known haemosporidian parasites are transmitted by blood-feeding dipteran insects [e.g.,1], biting Diptera remain the most plausible vectors.

However, nycteribiid flies frequently parasitize bat genera known to host *Nycteria* parasites, and the mitochondrial genome analysis recovered *Nycteria* and *Polychromophilus* as a monophyletic lineage; thus, a shared association with these highly specialized bat ectoparasites appears plausible. Overall, the common ancestor of *Nycteria* and *Polychromophilus* likely combined a loss of erythrocytic schizogony with a transition to bat-specific vectors, followed by diversification in tissue tropism and host associations [41–43]. Together, these patterns suggest that this putative bat-restricted clade represents not simply a host shift, but a broader reconfiguration of life-cycle strategy and transmission ecology within Haemosporida [8]. Our divergence-time estimates provide a robust temporal framework for understanding these evolutionary transitions.

Beyond life-cycle biology, *Polychromophilus* infects bats worldwide, occurring in both temperate and tropical regions, including the Americas, whereas *Nycteria* has so far been reported only from bats in Africa and Southeast Asia [48]. Their distributions overlap, particularly in Africa and Asia, where both parasite genera occur in local bat assemblages and infect bats from the families Rhinolophidae and Hipposideridae. However, their host associations differ markedly: *Polychromophilus* is most frequently associated with vespertilionid and miniopterid bats, whereas *Nycteria* primarily infects nycterid and rhinolophid hosts.

### *Nycteria-Polychromophilus* clade and molecular dating in Haemosporida

Calibrations are generally derived from fossil data or biogeographic events associated with the target organisms. In most groups, direct fossil evidence provides reliable temporal anchors. However, in the case of Haemosporida, fossil records are limited, and their interpretations remain controversial [49–51]. Consequently, molecular dating for this order requires secondary assumptions, such as reliance on the fossil record or the biogeographic history of their hosts [1,9]. A significant limitation of current models is that most available calibrations are restricted to mammalian *Plasmodium* [1,9]. This underscores the critical need for novel calibration points, particularly within lineages infecting Aves, Reptilia, and Chiroptera [1].

In the evolutionary history of Haemosporida, incorporating a calibration point for a putative *Nycteria*-*Polychromophilus* clade reduced the variance of divergence-time estimates. Although mitochondrial genome phylogeny is robust, it is not without limitations; mitochondrial genomes evolve at variable rates and under different selection regimes, likely influenced by haemosporidian vectors [9]. The inclusion of additional calibration constraints improves the precision of the time tree by reducing their highest probability densities (HPD). The constraints applied here—including the assumption of a monophyletic *Nycteria–Polychromophilus* group with an origin in early bats—improved the precision of the timetree by reducing their probability densities without producing major shifts in the overall evolutionary timescale.

While internal consistency among the calibration points does not prove their validity, it supports a reproducible temporal framework for testing evolutionary hypotheses [1]. The heterogeneity in haemosporidian mitochondrial substitution rates [9] increases uncertainty in divergence-time estimates, a problem that must be addressed with informative calibration priors. However, narrowly defined calibration priors were avoided, as all are derived from secondary sources in the absence of a haemosporidian fossil record. Imposing restrictive bounds may therefore bias posterior time estimates; thus, a degree of temporal uncertainty is unavoidable. Ultimately, expanding the molecular dataset beyond mtDNA may reduce sampling error and help mitigate the effects of rate heterogeneity, which are exacerbated by the small alignment size and lineage specific functional constraints [2,52].

Nevertheless, the most effective way to advance the field is to identify additional calibration constraints—ideally within genera such as *Haemoproteus* and *Leucocytozoon*—by discovering clades linked to clear biogeographic or host-speciation events. Simply increasing the number of extant species does not resolve the underlying issue; adding taxa without calibration can actually increase the variance in divergence-time estimates due to rate heterogeneity (Fig 6). Similarly, adding loci is only beneficial if they provide congruent phylogenetic signals, are not saturated, and exhibit consistent rate variation across lineages (e.g., avoiding genes under positive selection or those with highly divergent GC content) [1].

### Structural divergence within a bat-associated lineage

In addition to resolving topology, mitochondrial genomes provide structural information not accessible from short *cytb* gene fragments commonly used in haemosporidian diversity studies. Substantial mitochondrial gene-order rearrangements within a *Nycteria* lineage infecting *Nycteris* bats were confirmed here, as previously shown [21]. This is notable because mitochondrial genome architecture is otherwise highly conserved across haemosporidians [e.g., 9]. At present, the causes of these rearrangements remain unclear, but they may reflect prolonged independent evolution within chiropteran hosts. There is currently no evidence of functional consequences of this alternative gene order in parasite biology. Nevertheless, strong sequence-based monophyly and lineage-specific structural rearrangements underscore the evolutionary distinctiveness of this bat-associated clade.

## Materials and Methods

### Origin of samples and field sampling

All four samples analyzed in this study originated from previously published field studies conducted in Sierra Leone and Gabon (Table 4) [19,53]. The Sierra Leone sample (*Nycteria* sp., isolate NW2704) was obtained during an *ad hoc* bat survey conducted in the Gola Rainforest National Park (GRNP) at the border with Liberia, as part of a broader investigation of bat diversity and malaria parasite infections. Sampling permission was granted by the GRNP, with permits authorized by the Ministry of Agriculture and Forestry, Freetown, Sierra Leone. Sampling in Gabon was conducted with approval from the Centre National de la Recherche Scientifique et Technologique (CENAREST; permit AR0016/15/MESRS/CENAREST/CG/CST/CSAR). The Gabonese samples included two *Nycteria* isolates (NW3348, NW3236) and one *Polychromophilus* isolate (NW3328), collected during a rapid survey of haemosporidian infections in bats conducted at sites around Fougamou and in Waka National Park at the beginning of the dry season in June 2015. Bats were captured using standard canopy and ground-level mist nets placed along potential flyways and across different habitats, temporarily held individually in cotton bags, identified using established morphological keys [54,55], sampled, and released at the capture site. All capture and sampling procedures followed standard methods for monitoring mammal diversity and the guidelines of the American Society of Mammalogists [56].

**Table 4.**
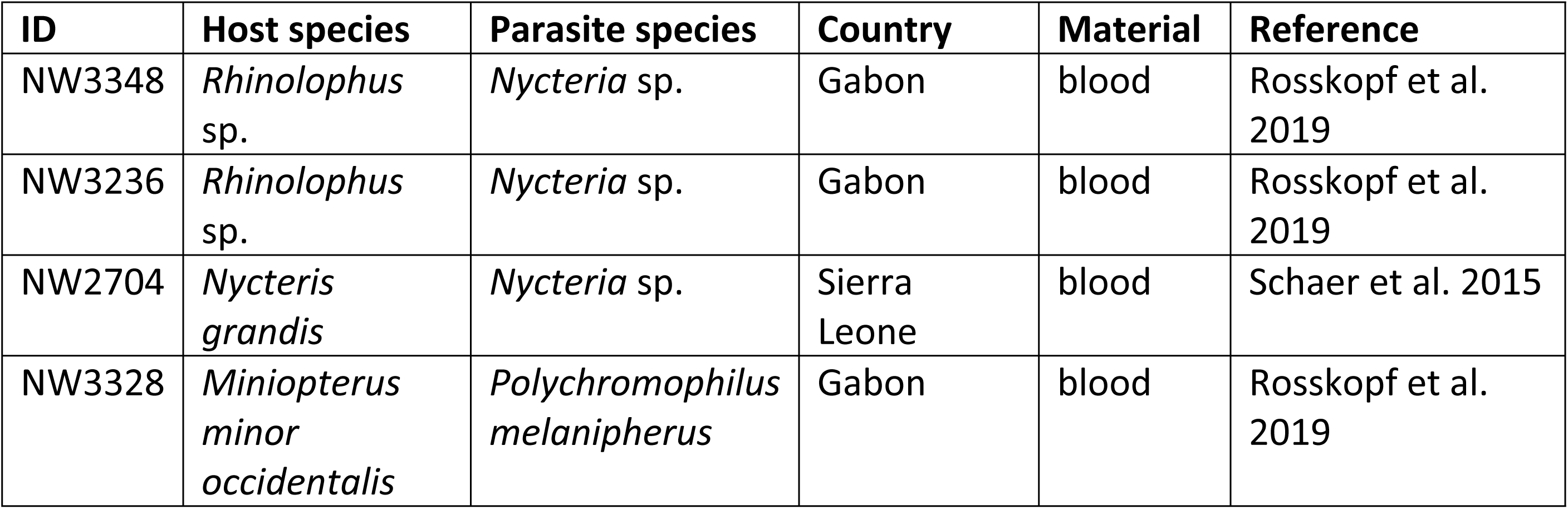
Samples for mitochondrial genome analysis.

### Blood collection, microscopy, and molecular screening

Blood samples (∼50 µl) were collected by venipuncture of the vein running along the anterior edge of the propatagium or antebrachial membrane, or the major vein in the interfemoral membrane, following Kurta and Kunz [57]. Immediately after collection, blood was used to prepare thin blood smears and to collect blood spots on DNA FTA cards (GE Healthcare) for morphological and molecular analyses. Thin blood smears were prepared in the field, fixed in 100% methanol, stained with Giemsa, and examined by light microscopy at 1,000× magnification with immersion oil to detect haemosporidian parasites.

Genomic DNA was extracted from dried blood spots on FTA cards using the DNeasy Blood & Tissue Kit (Qiagen, Hilden, Germany) following the manufacturer’s protocol for animal tissues with minor modifications. Blood spots were excised using sterile blades, and DNA was eluted in 80 µl of AE buffer. Initial molecular screening was performed using the AllTaq Master Mix Kit (Qiagen) with 2–4 µl of genomic DNA as template, targeting partial sequences of the mitochondrial cytochrome *b* gene (*cytb*) and the apicoplast caseinolytic protease (*clpC*), following previously described protocols [e.g. 13]. The resulting sequences and screening data were previously published [19,53]. For the selected samples (Table 4), selective whole-genome amplification was performed on extracted DNA using the REPLI-g Mini Kit (Qiagen, Hilden, Germany) according to the manufacturer’s instructions, followed by amplification and sequencing of complete mitochondrial genomes and additional nuclear loci.

### Mitochondrial genome amplification and sequencing

Mitochondrial genomes (≈6 kb) were amplified using LA Taq DNA Polymerase (TaKaRa, Shiga, Japan) with primers AE170 and AE171, following previously described protocols by Pacheco et al. [9,58]. Multiple independent PCR reactions were performed per sample to ensure reproducibility. PCR products were visualized by agarose gel electrophoresis, and amplicons of the expected size were excised under UV illumination and purified using the QIAquick Gel Extraction Kit (Qiagen, Hilden, Germany). DNA concentration and quality were assessed spectrophotometrically.

Purified mitochondrial PCR products were ligated into pGEM-T Easy vectors (Promega, Madison, WI, USA) following previously described protocols [9,58] and transformed into *Escherichia coli* JM109 competent cells by standard heat-shock. Recombinant clones were identified by blue–white screening on LB agar plates supplemented with ampicillin, IPTG, and X-Gal. White colonies were cultured in LB/ampicillin medium, and plasmid DNA was extracted using the innuPREP Plasmid Mini Kit (Analytik Jena). Successful insertion of mitochondrial fragments was confirmed by Sanger sequencing with M13 forward and reverse primers flanking the multiple cloning site [59].

For samples in which cloning was unsuccessful, PCR products amplifying the complete mitochondrial genome were purified directly using the NucleoSpin Kit (Macherey-Nagel) according to the manufacturer’s instructions. Both cloned inserts and purified non-cloned PCR products were sequenced by Sanger sequencing using overlapping internal primers spanning the entire mitochondrial genome. Sequencing was performed primarily in one direction per primer, with overlapping reads ensuring complete mitochondrial genome coverage (S1-S2 Tables).

### Assembly and annotation of mitochondrial genomes

Nucleotide sequences were imported into *GeneiousPrime* vs 2025.1.1 [60] for quality assessment, trimming, and manual editing. Complete mitochondrial genomes were assembled *de novo* and subsequently validated by mapping reads against haemosporidian reference mitochondrial genomes obtained from NCBI (*Nycteria medusiformis*: KX090645, *Nycteria gabonensis*: KX090647; *Polychromophilus murinus*: OP380907). Gene annotations were transferred from reference sequences and verified by manual inspection of open reading frames and translation accuracy for protein-coding genes, *cytB, cox1,* and *cox3*. The complete mtDNA genome sequences were submitted to GenBank under the accession number PZ224672-PZ224675.

### Amplification and selection of additional nuclear markers

Although the principal inferences are based on near-complete mitochondrial genomes, additional nuclear loci were examined to assess whether similar relationships are recovered across independent genetic markers. Selected nuclear markers were amplified for two representative samples, NW3348 (*Nycteria* sp.) and NW3328 (*Polychromophilus* sp.), following the gene set and laboratory protocols described by Galen et al. [10]. Because previous studies utilizing 21 nuclear loci suffered from uneven taxon representation, with *Polychromophilus* and *Nycteria* particularly underrepresented, we targeted a subset of nine markers to ensure complete representation for our target genera and improve overall data comparability.

The amplified loci comprised partial sequences of the following markers: metabolite/drug transporter (DrugT), splicing factor 3B subunit 1 (SF3B1), pre-mRNA-processing factor 6 (PRPF6), DNA polymerase delta subunit 1 (POLD1), DNA replication licensing factor MCM5, importin beta (KPNB1), protein transport protein SEC24A, eukaryotic translation initiation factor 2 gamma subunit (*EIF2*), and the LCCL domain-containing protein (CCP1). All nuclear loci were amplified using a nested PCR approach, with inner primers incorporating either a CAG (5′-CAGTCGGGCGTCATCA-3′) or M13R (5′-GGAAACAGCTATGACCAT-3′) sequencing tag. PCR amplifications were performed using the ExTaq DNA Polymerase Kit (TaKaRa, Shiga, Japan). PCR products were purified using the NucleoSpin Kit (Macherey-Nagel) according to the manufacturer’s instructions and sequenced by Sanger sequencing using CAG- or M13-tagged inner primers. Nuclear gene sequences were submitted to the GenBank under the accession number PZ035787 – PZ035798.

### Phylogenetic analyses

#### mtDNA genomes

Two nucleotide alignments were performed with ClustalX v2.0.12 and Muscle, as implemented in SeaView v4.3.5 [61], and manually edited. The first alignment (5,052 bp excluding gaps) comprised the new sequences generated here (*Nycteria* sp. NW3348 and *Polychromophilus* sp. NW3328) and 157 mtDNA sequences from seven Haemosporida genera: *Leucocytozoon*, *Haemoproteus*, *Haemocystidium*, *Plasmodium*, *Hepatocystis*, *Polychromophilus*, and *Nycteria* available in the GenBank. The alignment was annotated according to the gene models proposed by Feagin et al. [20], identifying fragmented small subunit ribosomal RNAs (SSU rRNA), large subunit ribosomal RNAs (LSU rRNA), and the three protein-coding genes. Protein-coding genes (*cox1*, *cox3*, and *cytb*) were identified based on conserved open reading frames and alignment with reference sequences. Gene boundaries were refined through comparative alignment across taxa to ensure consistency in annotation. All sequences were manually inspected to verify gene structure and to detect potential sequencing or assembly errors. Then, this alignment was divided into six partitions corresponding to the three nonprotein-coding regions between the ORFs (fragmented SSU rRNA and LSU rRNA) and the three protein-coding genes.

A second alignment was constructed, consisting of 162 sequences (3,270 bp, excluding gaps), restricted to the three protein-coding genes (*cox1*, *cox3*, and *cytb*). This dataset included 162 mtDNA sequences, including the three *Nycteria* sequences with rearranged mitochondrial genomes: *Nycteria medusiformis* (KX090645, [21]), *Nycteria* sp. (KX090646, [21]), and the new sequence for *Nycteria* sp. (NW2704) obtained in this study. This alignment was divided into three partitions, one for each protein-coding gene, to account for differences in substitution rates and models among genes.

Phylogenetic hypotheses were explored based on the first (near-complete mtDNA genome) and second (only CDS) alignments using Bayesian Inference (BI) implemented in MrBayes v3.2.7 with the default priors [22] and a maximum-likelihood (ML) method implemented in IQ-TREE v2.3.1 [23]. A general time reversible model with gamma-distributed substitution rates and a proportion of invariant sites (GTR + Γ +I) was selected based on the lowest Bayesian Information Criterion (BIC) scores as estimated in MEGA v12.1 [62]. For the BI analysis, posterior probabilities for the nodes were estimated using two independent Markov Chain Monte Carlo (MCMC) runs of 6 × 10⁶ generations, sampling every 500 steps. Convergence was confirmed by a potential scale reduction factor (PSRF) between 1.00 and 1.02, effective sample sizes (ESS) > 200, and an average standard deviation of split frequencies < 0.01 [22]. A 25% “burn-in” was discarded for each run. For the ML analysis, support values were assessed via Ultrafast Bootstrap approximation (UFBoot) with 1,000 replicates. Phylogenetic trees were visualized using FigTree v1.4.4 (http://tree.bio.ed.ac.uk/software/figtree/). All parasite species names and GenBank accession numbers used in both analyses are provided within the figures.

#### Nuclear genes

All nuclear DNA sequences were assembled, quality-checked, and manually curated using *GeneiousPrime* vs 2025.1.1 (Biomatters), and aligned with reference sequences primarily derived from Galen et al. [10] as well as additional taxa retrieved from NCBI using the MAFFT alignment algorithm. Each nuclear locus was first analyzed separately to evaluate gene-specific phylogenetic signal and potential discordance among nuclear markers. Subsequently, individual gene alignments were concatenated for combined phylogenetic inference. Seven taxa with substantial missing data (i.e., represented by only one to three loci) were removed from the concatenated alignment prior to phylogenetic analyses, while being retained in single-gene alignments. The final concatenated dataset comprised 7,377 bp, of which 6,453 were gap-free.

Phylogenetic relationships for the nuclear data were inferred for each individual gene alignment and for the concatenated dataset using a Bayesian inference (BI) and maximum likelihood (ML) approaches. For all analyses, datasets were partitioned by codon position, with first, second, and third codon positions specified separately for all protein-coding loci. For the concatenated analyses, datasets were partitioned by gene and by codon position. Optimal partitioning schemes and nucleotide substitution models for the Bayesian analyses were identified using PartitionFinder v2 based on BIC, and the selected models were applied in subsequent Bayesian analyses (see S3 Table).

Bayesian inference was conducted in MrBayes v3.2.7 via the CIPRES Science Gateway using two independent runs with four Markov chains each (one cold and three heated). Temperature, number of generations, and substitution models used for the analyses are summarized in S3 Table S3. Convergence was assessed using potential scale reduction factors (PSRFs) between 1.00 and 1.02 and an average split-frequency standard deviation below 0.01; all parameters achieved effective sample sizes (ESS) greater than 1,000. The sampling frequency was 1,000, and the first 25% of sampled trees were discarded as a burn-in.

Maximum likelihood analyses were performed using IQ-TREE v2.3.1. The concatenated dataset was partitioned by gene and codon position, and the best-fit substitution models and optimal partitioning scheme were selected using ModelFinder according to the Bayesian Information Criterion (BIC). To avoid over-partitioning, similar partitions were allowed to merge during model selection. Nodal support was assessed using the ultrafast bootstrap approximation (UFBoot) with 1,000 replicates. All resulting phylogenetic trees were visualized and edited using FigTree v1.4.4. Individual gene trees are provided in the Supplementary Material (S1 Fig).

### Divergence times and rate of evolution of Haemosporida using mitochondrial genome

Divergence times were estimated based on the near-complete mitochondrial genome alignment (159 mtDNA sequences, 5,052 bp excluding gaps) using both a Bayesian approach implemented in MCMCTree [27–29] and a non-Bayesian framework based on the RelTime method [30] implemented in MEGA v12.1 [62]. Unlike Bayesian methods, RelTime does not assume an explicit statistical model for rate variation among lineages [30]. However, the method is not assumption-free, as it computes the rate of an ancestral branch as the average of its two immediate descendant branches [30]. Rate autocorrelation was assessed using CorrTest [31], also implemented in MEGA v12.1.

In MCMCTree, divergence times were estimated using the approximate likelihood method [63] under the GTR+Γ substitution model, based on the MrBayes topology derived from the mitochondrial genome alignment. A prior for the overall rate parameter was specified following the guidelines in the MCMCTree manual, based on a baseml analysis conducted under a strict (global) clock model with a point calibration (calibration 2 at 29.22 Ma; see below). The birth-death process parameters were set to “BDparas= 1 1 0”. Both independent rates [64] and autocorrelated rates models [28,65] were explored. Calibration priors were specified as uniform probability distributions with soft bounds (see below). To ensure convergence, two independent MCMC runs were performed for each combination of rate model and calibration scheme until ESS values exceeded 200 after discarding the burn-in.

RelTime analyses were performed using the command-line version of MEGA v12.1 [62]. The substitution model was the same as that used in Bayesian analyses (GTR+Γ). The same minimum and/or maximum bounds applied in the Bayesian framework were also used. To evaluate the influence of calibrations on divergence time estimates, the protocol described by Battistuzzi et al. [34] was implemented. RelTime was also utilized to estimate branch-specific relative evolutionary rates in Haemosporida, which were visualized using FigTree v1.4.4 (http://tree.bio.ed.ac.uk/software/figtree/).

For both the BI and non-BI methods, three distinct calibration scenarios were explored using uniform priors, so that no specific time points were favored within the defined intervals. Calibration constraints incorporated some host fossil data from the Paleobiology Database (http://www.paleodb.org/, accessed October 2025) and a combination of molecular dating of lemurs [17] utilizing both recent and ancient fossils [66] and divergence estimates from the TimeTree database (https://www.timetree.org/, accessed October 2025). While these represent secondary calibrations, they remain the most robust temporal constraints available, given the absence of a reliable haemosporidian fossil record [1].

The first calibration scenario comprised four calibration constraints previously tested by [9,16,17]. (1) The divergence between Papio-Macaca parasites was assigned uniform prior with soft bounds of 6.0–14.2 Ma based on the cercopithecine fossil record [67,68]; (2) the Human-Macaca split was calibrated with soft bounds of 24.44–34.0 Ma, corresponding to the hominoid–cercopithecoid divergence; (3) the origin of Lemuridae–Indriidae clade, with soft bounds of 20.0–42.0 Ma (mean age = 34.83 Ma; 95% CrI = 27.18–42.25 Ma [9,66]); and (4) the Bovinae–Antilopinae divergence, with soft bounds of 16.0–28.1 Ma [9,69]. This scenario yielded divergence estimates for *Polychromophilus* and *Nycteria* that were consistent with the inferred evolutionary origins of their chiropteran hosts [32,33]. Therefore, two additional calibration scenarios were explored.

In the second calibration scenario, a 40–65 Ma uniform prior was added for the origin of chiropteran parasites, in addition to the four initial calibration priors. The lower bound (40 Ma) reflects the minimum divergence age of most extant chiropteran families, whereas the upper bound (65 Ma) incorporates molecular clock estimates suggesting that the crown-group of bats originated near the Cretaceous–Paleogene (K–Pg) boundary, approximately 62–65 Ma [32,33]. The third calibration scenario included these five calibration priors, along with an additional 65–70 Ma uniform prior placed at the divergence between bats and other mammalian lineages, estimated to have occurred approximately 65–70 Ma, shortly after the mass extinction of non-avian dinosaurs [32,70–74].

In MCMCTree, a calibration prior must be specified for the root of the tree. For all scenarios explored, the origin of Palaeognathae (56.8–86.8 Ma, [69]) was used as the root calibration prior. The lower bound (56.8 Ma) corresponds to the fossil *Lithornis celetius* [75] from the Fort Union Formation (early Tiffanian) of Montana and Wyoming, whereas the upper bound (86.8 Ma) reflects a soft maximum based on the age of the Niobrara Chalk Formation, dated as Santonian (86.3–83.6 ± 0.5 Ma). As with the other calibrations, a uniform prior with soft bounds was used. The root calibration was not included in the RelTime analyses because this method does not accommodate calibrations at the root node, as the rate estimation algorithm assumes equal evolutionary rates between the ingroup and outgroup. In MCMCTree, divergence times were estimated using the approximate likelihood method [63].

## Acknowledgments

We thank the Gola Rainforest National Park (GRNP) and the Ministry of Agriculture and Forestry in Sierra Leone for providing the necessary sampling permits and field support. We are also grateful to the Centre National de la Recherche Scientifique et Technologique (CENAREST) in Gabon for authorizing the research activities conducted in Fougamou and Waka National Park. We thank Kai Matuschewski, Natalie Weber, Annika Hillers, Jerry Garteh, Amadu Jusu, Brima S. Turay, Nadia Wauquier, Jana Held, Markus Gmeiner, and Isabella Eckerle for their contributions to sample collection and for their work in the original projects in which the samples were obtained.

M. Andreína Pacheco and Ananias A. Escalante are supported by the US National Science Foundation (Grant No. NSF-DEB 2146653). Juliane Schaer is supported by the German Science Foundation (grant number 437846632). Beatriz Mello is supported by Fundação Carlos Chagas Filho de Amparo à Pesquisa do Estado do Rio de Janeiro (FAPERJ) (Grants No. E-26/201.446/2022 and E-26/204.534/2025) and by Conselho Nacional de Desenvolvimento Científico e Tecnológico (CNPq) (Grants No. 311231/2022-5). The founders had no role in study design, data collection and analysis, decision to publish, or preparation of the manuscript.

## Data accessibility

All sequence data generated in this study, including the complete mitochondrial genomes for *Nycteria* sp. and *Polychromophilus* sp., have been deposited in the GenBank database under the accession numbers PZ224672-PZ224675 (mtDNA) and PZ035787 – PZ035798 (nuclear gene sequences).

## Supporting information captions

**S1 Fig. Bayesian and Maximum Likelihood phylogenetic hypothesis of haemosporidian species based on nine nuclear genes.** The nodes’ values are posterior probabilities, and bootstrap support is reported as a percentage. GenBank accession numbers are shown. *Nycteria* and *Polychromophilus* parasites are highlighted in blue and red colors respectively. (A) *Sec24A* - Red asterisks indicate inconsistencies between both phylogenetic methods (BI and ML). (B) *DrugT* - Red asterisks indicate inconsistencies between both phylogenetic methods (BI and ML). (C1) *POLD1 –* BI analysis. (C2) *POLD1 –* ML analysis. (D1) *Ccp1 –* BI analysis. (D2) *Ccp1 –* ML analysis. (E1) *KPNB1 –* BI analysis. (E2) *KPNB1 –* ML analysis. (F1) *Sf3b1 –* BI analysis. (F2) *Sf3b1 –* ML analysis. (G1) *Prpf6 –* BI analysis. (G2) *Prpf6 –* ML analysis. (I1) *MCM5 –* BI analysis. (I2) *MCM5 –*ML analysis.

**S2 Fig. (A) Comparation of divergence time estimates using autocorrelated versus the independent rate models in MCMCTree under three scenarios. (B). Comparation of divergence time estimates using autocorrelated versus the independent rate models in MCMCTree and RelTime under three scenarios.** See Materials and Methods for details.

**S3 Fig. Relative times without calibrations versus absolute times with calibrations (Ma) as estimated using RelTime in MEGA v under three scenarios (see Materials and Methods).**

**S1Table. Primers used for mitochondrial genome amplification and core sequencing**

**S2 Table. Internal mitochondrial primers designed in this study.**

**S3 Table. Information for Bayesian inference analyses of nuclear datasets.**

